# Diverse somatic Transformer and sex chromosome karyotype pathways regulate gene expression in Drosophila gonad development

**DOI:** 10.1101/2024.08.12.607556

**Authors:** Sharvani Mahadevaraju, Soumitra Pal, Pradeep Bhaskar, Brennan D. McDonald, Leif Benner, Luca Denti, Davide Cozzi, Paola Bonizzoni, Teresa M. Przytycka, Brian Oliver

## Abstract

The somatic sex determination gene *transformer* (*tra*) is required for the highly sexually dimorphic development of most somatic cells, including those of the gonads. In addition, somatic *tra* is required for the germline development even though it is not required for sex determination within germ cells. Germ cell autonomous gene expression is also necessary for their sex determination. To understand the interplay between these signals, we compared the phenotype and gene expression of larval wild-type gonads and the sex-transformed *tra* gonads. XX larval ovaries transformed into testes were dramatically smaller than wild-type, with significant reductions in germ cell number, likely due to altered geometry of the stem cell niche. Additionally, there was a defect in progression into spermatocyte stages. XY larval testes transformed into ovaries had excessive germ cells, possibly due to the earlier onset of cell division. We suggest that germ cells are neither fully female nor male following somatic sex transformation, with certain pathways characteristic of each sex expressed in *tra* mutants. We found multiple patterns of somatic and germline gene expression control exclusively due to *tra*, exclusively due to sex chromosome karyotype, but usually due to a combination of these factors showing *tra* and sex chromosome karyotype pathways regulate gene expression during Drosophila gonad development.

## INTRODUCTION

The somatic sex determination pathway in *Drosophila melanogaster* is a textbook example of an RNA splicing cascade. Somatic cells use a sex-specific X chromosome karyotype to initiate the cascade. XX flies are female and single X flies are male (Calvin B. Bridges 1914; Cline and Meyer 1996; Salz and Erickson 2010). The Y chromosome is essential for spermatogenesis but plays no role in somatic sex differentiation (C. B. Bridges 1916; Hafezi et al. 2023). Two X chromosomes (XX) activate the *Sex lethal* (*Sxl*) locus, which encodes an RNA-binding protein that autoregulates *Sxl* pre-mRNA to maintain female identity (Bell et al. 1988; Salz and Erickson 2010). In addition, Sxl protein regulates *transformer* (*tra*) pre-mRNA splicing to produce a long open reading frame leading to functional Tra protein in XX cells, while a truncated non-essential peptide is produced in X(Y) cells (Inoue et al. 1990). Tra is an RNA-binding protein that in partnership with *transformer* 2 (*tra 2*) directly splices *doublesex* (*dsx*) and *fruitless* (*fru*) transcription factors that regulate almost all sexually dimorphic development and behavior in flies (Ryner et al. 1996; Camara, Whitworth, and Van Doren 2008). There are only a few other known direct targets of *tra* (Hudry, Khadayate, and Miguel-Aliaga 2016). The Tra protein along with the cofactor protein coded by *tra 2* locus (non-sex specific) (Amrein, Gorman, and Nöthiger 1988), directly binds to a splicing enhancer and alternatively splices *dsx* pre-mRNA (Belote and Baker 1982; Baker and Wolfner 1988) to produce a female specific isoform, DSX^F^, which switches development to female. In males, a single X leads to the absence of female-specific Sxl and Tra, which leads to a male specific *dsx* mRNA that encodes DSX^M^, which switches development to male **(Fig. 1A)**. The somatic sex determination gene cascade also controls the sex of the germline in addition to the germline autonomous regulation (Illmensee and Mahowald 1974; Steinmann-Zwicky, Schmid, and Nöthiger 1989; Pauli and Mahowald 1990; Steinmann-Zwicky 1992; B. Oliver, Kim, and Baker 1993; Brian Oliver 2002). Similar to somatic cells, the sex of the germline is determined by a sex-specific X chromosome karyotype (XX is female germ cell and single X is male germ cell) (Schüpbach 1985; Brian Oliver 2002). Importantly, the germ cell transplantation experiments suggest that the sex of somatic and germline components must match for correct sex-specific differentiation of germ cells and functional gamete production (Schüpbach 1985; Brian Oliver 2002; Gilboa and Lehmann 2004a).

**Fig 1.**
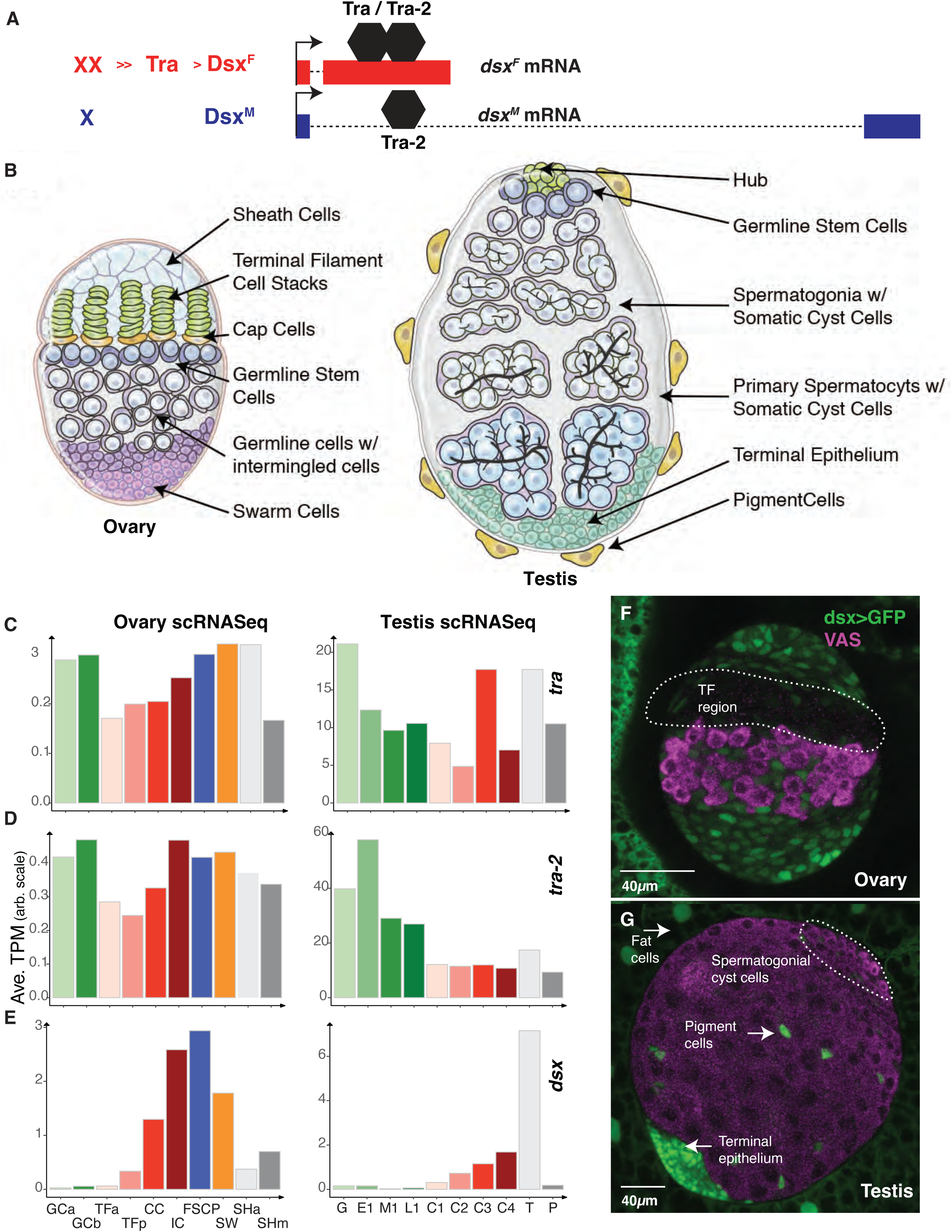
Sex Determination gene expression in Drosophila larval gonads at single cell resolution. A. Model of initial somatic sex determination signals. The karyotype in female (XX) versus male (X, also has a Y, where Y is not a part of sex determination signal but involved in fertility). In response to XX karyotype, Tra protein is produced and binds to *dsx* mRNA along with Tra2 protein leading to the production of DsxF isoform. In response to single X in the absence of Tra protein, leads to the production of DsxM isoform. B. Cartoons of third instar larval ovary and testis. Ovary representing sheath cells, terminal filaments, cap cells, germline stem cells, germline cells with intermingled cells and swarm cells. Testis representing hub, germline stem cells, spermatogonia with somatic cyst cells, primary spermatocytes, terminal epithelial cells, and pigment cells. C., D., and E. are the gene expression (Average Transcript per Million (TPM) from scRNASeq) of *tra*, *tra*-2 and *dsx*, in different cell types of larval ovary and testis. F. and G. Immunostained gonads where dsx-Gal4 is driving uas-GFP expression in ovary and testis. White arrows representing Dsx expressing cells. The TF region in the ovary is the Terminal Filament region represented by the dotted line). Gal4-dsx > uas-GFP: Green (fatbody, sheath cells, intermingled cells, terminal epithelium), VAS: Magenta (germ cells). Scale bars 40 *μ*m

The sex determination cascade genes are required in the somatic cells of the gonads that drive the highly sexually dimorphic gonad development. Mutations in these genes in the soma cause clear gonadal transformation phenotypes: the development of testes in XX; *tra* loss of function flies *(tra* null) and the development of ovaries in XY; *tra* gain of function flies (expressing female *tra* cDNAs) (Sturtevant 1945; McKeown, Belote, and Boggs 1988; Steinmann-Zwicky 1994; Brian Oliver 2002; Evans and Cline 2007). Sex-transformed gonads are clearly defective at eclosion and show defects not only in the somatic cells but also gross defects in the germline, even-though clonal analysis of these genes shows that they are not required for sex determination function within the germ cells (Marsh and Wieschaus 1978; Schüpbach 1982; 1985; Pauli and Mahowald 1990; B. Oliver, Kim, and Baker 1993; Horabin et al. 1995; Brian Oliver 2002; Murray, Yang, and Van Doren 2010).

An important characteristic of sex-transformed adult gonads is a highly variable germline phenotype, and some studies reported the germ cells as female, some as males, some others as neither or both (Brown and King 1961; Seidel 1963; Nöthiger et al. 1989; Casper and Van Doren 2006). XX flies transformed from females to males have testes with very few adult germ cells, while XY flies transformed from males into females have ovarian tumors composed of large numbers of germ cells that fail to differentiate (Steinmann-Zwicky 1994). The germ cell loss in XX females transformed into males is more pronounced as those flies age, so most of what we know about the phenotype of these germ cells is from the rare surviving germ cells. This has limited our understanding of the underlying mechanism. As the wild-type ovaries and testes in larvae are overtly dimorphic **(Fig. 1B)** but do not have differentiated gametes, we wondered if sex transformations might be appropriate to decipher at this stage.

The larval gonads at the end of larval development are ellipsoids and embedded in a sheet of fatbody cells (Rosales-Nieves et al. 2023). When viewed with the anterior and posterior poles vertically, ovaries have an equatorial band of germ cells (Rongo et al. 1997). These germ cells are at or near the end of a post-embryonic mitotically quiescent period (Lebedeva et al. 2018). The future ovarian sheath cells and the somatic cells of the female germline niche (terminal filament and cap cells) are located anterior to the germ cells. These cells are also beginning to arrange into the adult niche configuration (Drummond-Barbosa 2019). Germline stem cells are located nearest the niche, while more posterior germ cells might be destined to differentiate directly into egg chambers. Female germ cells are fully surrounded by somatic intermingled cells (Gilboa and Lehmann 2006; Banisch et al. 2021). The germ cells take up the bulk of the larval testis. The male germ cell divisions begin in embryogenesis shortly after gonad formation and continue into larval stages (Wawersik et al. 2005). By the third instar, there are self-renewing germline stem cells at the niche, defined by a tight group of cells called the hub (Hardy et al. 1979; Drummond-Barbosa 2019). Differentiating daughter germ cells undergo four rounds of mitosis in interconnected cysts and then enter a prolonged premeiotic G2 phase. These primary spermatocytes are characterized by nuclear and cytoplasmic growth and the activation of a complex gene expression program required for post-meiotic spermiogenesis (Hennig 1996). Like in the ovary, the male germ cells are fully surrounded by somatic cyst cells. We compared these wild-type larval gonads to the sex-transformed gonads developed by *tra* perturbation in the somatic cells of the gonads. We identified specific cell types in the gonads by immunostaining and transcriptomics approaches (RNA-seq, PacBio long-read, and single-cell sequencing) that revealed different pathways sex determination and germline differentiation are regulated in gonads.

## RESULTS

### Somatic sex determination gene expression in third instar larval gonads

Even though we know the function of somatic sex determination cascade genes genetically, we know very little about their expression in larval gonads especially in specific cell types. Single cell expression profiles exist for both wild-type larval ovaries and testes. At published resolutions (Slaidina et al. 2020; Mahadevaraju et al. 2021), 10 different cell types exist in each gonad that provide a chance to examine sex determination and differentiation gene expression in specific cell types. Somewhat surprisingly given the lack of a germline requirement, *tra* was highly expressed in the germ cells of both sexes relative to expression in the somatic cells (**Fig. 1C**). Read analysis indicated that at least some of the Tra mRNAs encode the female-specific isoform (**Table S6**). Expression of the Tra cofactor, *tra-2,* showed a similar expression profile among gonadal cell types (**Fig. 1D**). There is no known role for Tra in the germline, but the fact that it is expressed in germ cells means that it may play a role, albeit not one required for fertility.

The *dsx* mRNAs were expressed at high levels in somatic gonad cells, with the notable exception of the cell clusters that give rise to the terminal filaments (TFa), which are somatic components of the germline stem cell niche **(Fig. 1E)**. In larval testes, *dsx* mRNA is poorly expressed in all the germ cells (spermatogonia and spermatocytes), with very high expression in the terminal epithelium and lower expression in the somatic cyst cells. The *dsx* locus is expressed in all the somatic cells of the newly coalesced embryonic gonads (Hempel and Oliver 2007), but not in the germ cells, raising the possibility that *dsx* is transient in some gonad cell types. In larval gonads, the *dsx* expression pattern appears to be entirely somatic cell type-specific, suggesting that any role Tra and Tra-2 play in the germline is Dsx independent.

We also analyzed the expression of a dsx-Gal4 driver (Robinett et al. 2010; Rideout et al. 2010) which was clearly expressed in the subsets of somatic gonadal cells. We saw driver expression in all ovarian cell types except the germ cells and the somatic cells just anterior to the germ cells, where the somatic niche is forming (**Fig. 1F**). The terminal filament stacks were unstained in these experiments, which is entirely consistent with the scRNA-seq data for *dsx* mRNAs. In the testes, we observed dramatic expression of the *dsx* driver in the terminal epithelium, again as predicted by the single-cell expression analysis (**Fig. 1G**). We saw weak, but consistent, expression of the *dsx* driver in the somatic cyst cells surrounding the spermatogonia, but not in the other somatic cyst cells. We also observed the highest *dsx* RNA in the somatic cyst cell cluster C4, suggesting that this cluster represented the cyst cell surrounding the spermatogonia. Thus, any *tra* effects on many of the somatic cells could well be mediated by *dsx*.

### The *transformer* sex-transformation phenotypes

We examined the morphology of four types of gonads: wild-type ovaries and testes, testes transformed into ovaries by ectopically expressing Tra using an allele that rescues *tra* and ovaries transformed into testes by the absence of *tra* using a molecular null **(Table S1)**. If we successfully removed or over expressed *tra*, then *dsx* expression should be altered. When we examined *dsx* expression in these gonads by RNA-seq, we observed female splicing in the XX ovaries and the XY testes transformed into ovaries, and male splicing in the XY testes and the XX ovaries transformed into testes, indicative of a molecular-level sex transformation (**Fig. 2A**).

**Fig 2.**
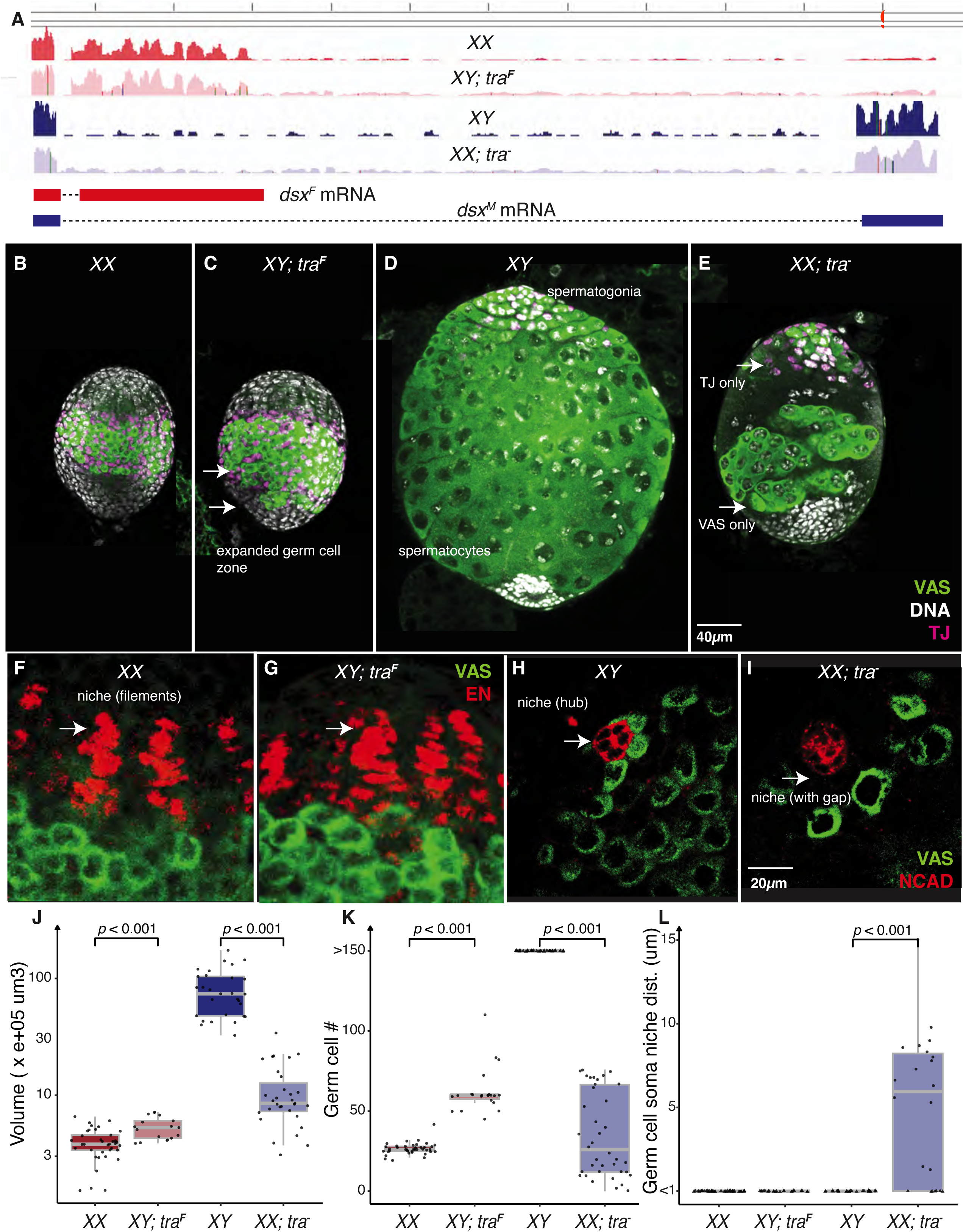
Sex transformed Drosophila larval gonads. A. Visualizing the gene expression (reads) of *dsx* locus on the Genome browser from the RNA-seq data of larval gonads from wild type ovary (XX), testis sex transformed into ovary (XY; *tra*^F^), wild type testis (XY) and ovary sex transformed into testis (XX; *tra*^-^). The dsx^F^ mRNA is represented by Red and dsx^M^ mRNA is represented by blue genomic tracks to follow female and male specific splicing patterns. XX; *tra*^-^ (Df(tra)/tra^1^) is a progeny from w^1118^; Df(3L)st-j7, Ki/TM6 females and w^1118^/Y; tra[1]/TM6 males. XY; *tra*^F^ (U2af-*tra*^F^/+; *tra*^5^) is a progeny from X/X; tra[F.U2af50]/+; tra[5] females and FM7, act-gfp/Y males. B, C, D and E are Immuno stained XX, XY; *tra*^F^, XY and XX; *tra*^-^ gonads to observe the morphology VAS: green (Germ cells), TJ: Magenta (cyst cells), DAPI: gray (Nuclei). Scale bar 40 *μ*m F, G, H and I are the Immuno stained XX, XY; *tra*^F^, XY and XX; *tra*^-^ gonads to observe Niche regions VAS: Green (Germ cells), Engrailed: Red (terminal filaments), Ncad: Red (hub), Scale bars 20 *μ*m White arrows in F and G are pointing to similar looking niche regions in XX and XY; *tra*^F^. White arrow in H is pointing no-gap between the hub to the nearest germ cells. White arrow in I is pointing the gap between the hub to the nearest germ cells. J. Volume (V=4/3!abc) of XX, XY; *tra*^F^, XY and XX; *tra*^-^ larval gonads. K. Total number of germ cells in the larval gonads of the genotype represented. L. The distance between the somatic cells in the niche region to the nearest germ cell. Two-sample Wilcoxon Rank Sum (Mann-Whitney) test is utilized to find the statistical significance in J - L. In L, triangles indicate numbers less than 1 were raised to 1.

Even at this larval stage, there were clear differences among the wild-type and sex-transformed gonad morphologies. We used Vasa (Vas) (Hay, Jan, and Jan 1990) to mark germcells and Traffic jam (Tj) (M. Li et al. 2019) to mark closely associated somatic support cells. XX ovaries (**Fig. 2B**) were characterized by an equatorial band of Vas^+^ germ cells and closely associated somatic intermingled cells that were Tj^+^. The XY testes transformed into ovaries by the action of Tra (**Fig. 2C**) were grossly similar but had a more posteriorly spread germ cell region. Like the wild-type ovaries, the testes transformed into ovaries had Tj^+^ cells in close association with the germ cells. These XY testes transformed into ovaries had a significantly higher estimated volume and more germ cells than XX ovaries (**Table S2, Fig. 2J, K**).

The wild-type XY testes (**Fig. 2D**) had spermatogonial cysts at the apex and 16-cell primary spermatocyte cysts through most of the rest of the gonad. The XX ovaries transformed into testes were dramatically smaller than wild-type, with heavy reductions in germ cell number (**Fig. 2E, J, K**). Probable spermatogonia were often seen at the apex, while enlarged germ cell cysts (defective spermatocytes?) were seen more posteriorly (**Fig. 2E**). Tj^+^ cells were sometimes seen in tight association with the germ cells, this was not always the case. This larval phenotype was highly variable. Uniquely, among the four gonad types, we observed occasional germ cells without associated Tj^+^ cells and clusters of Tj^+^ cells without germ cells in ovaries transformed into testes (**Fig. 2E**). This striking lack of somatic support cell / germ cell contact could help explain the severity of the XX testis phenotype.

The germ cell niche regulates germ cell number in both sexes. Given the excess germ cell numbers in XY testes transformed into ovaries and the paucity of germ cells in XX ovaries transformed into testes, we carefully examined the germ cell niche regions. In third instar XX ovaries Engrailed-positive (En^+^) somatic cells (Gilboa 2015) intercalated to arrange into terminal filaments (**Fig. 2F**), an important somatic cell to promote germline stem cell renewal and daughters to develop into eggs. The posterior most ends of these filament stacks cells were in contact with the anterior groups of germ cells (**Fig. 2L**). We did not detect overt differences in niche structure in the XY testes transformed into ovaries (**Fig. 2G, L**). In XY testes, germline stem cells were arranged (**Fig. 2H**) around Neural Cadherin positive (Ncad^+^) hub cells (Greenspan, de Cuevas, and Matunis 2015). Like the terminal filaments in ovaries, the hub is essential for maintaining the germline stem cell population in testes. In stark contrast to the wild-type situation, XX ovaries transformed into testes had germ cells that were often detached from the hub (**Fig. 2I, L**). This is an important observation, as stem cell renewal depends on contact between the germ cells and the hub. In the adult, failed contact between germ cells and the hub eventually results in full depletion of the germline (Tulina and Matunis 2001). This altered geometry of the niche immediately suggests that stem cell depletion contributes to the loss of germ cells in XX females transformed into males.

### Gene expression in sex-transformed gonads

To investigate gene expression changes due to somatic sex transformation in gonads, we developed RNA-Seq gene expression profiles for these four gonad types in quadruplicate (see Methods). We used Salmon (Patro et al. 2017) for expression quantification and interpreted the results using ontologies (Klopfenstein et al. 2018) and expression profiles from larval ovary and testes single cell atlases (Slaidina et al. 2020; Mahadevaraju et al. 2021).

Our experiment had two variables, so to incorporate both somatic sex and X chromosome karyotype, we clustered all the genes showing differential gene expression after performing a two-way analysis of variance (ANOVA, **Fig. 3A**, **Table S3**, see Methods). We found groups of genes that were highly expressed in both XX and XY ovaries and not in testes, as well as genes expressed in XY and XX testes and not in ovaries. These genes are candidates for direct or indirect regulation by Tra. We observed a smaller set of genes where the expression followed X chromosome numbers. These genes could be regulated upstream of *tra* in the soma, or perhaps are expressed in the germline where they follow the germline XX signal. However, it was immediately clear that most genes were directly or indirectly controlled by both the presence or absence of Tra and the number of X chromosomes (**Fig. 3A**). We also saw genes highly expressed in only one gonad type, especially wild-type testes. The large cluster of genes expressed highly only in XY testes contained many genes known to be expressed in primary spermatocytes in bulk, and single-cell studies (Vedelek et al. 2018; Witt et al. 2019; Raz et al. 2023). These results strongly suggest that XY testes transformed into ovaries do not have germ cells that reach the spermatocyte stage. Similarly, XX ovaries transformed into testes might have few to no germ cells that reached the primary spermatocyte stage, even though they did appear to have cysts with enlarged cells, at least superficially similar to spermatocytes. We will return to the depletion of aborted spermatocytes later in the manuscript. These cases of gene expression in only one of the four gonad genotypes indicates that many genes require the coordinated action of Tra and ill-defined X chromosome-dependent factors.

**Fig 3.**
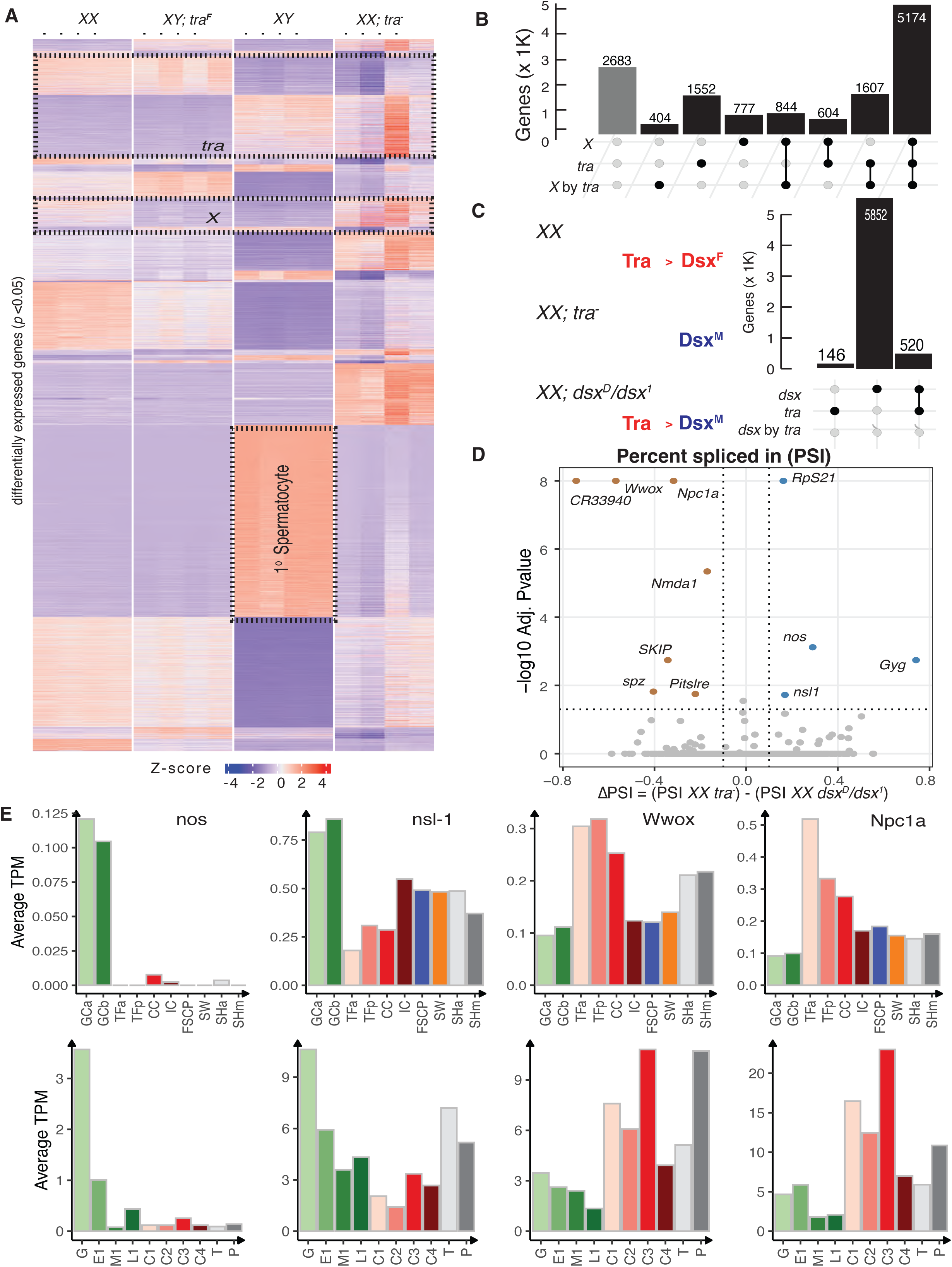
Transcriptomics analysis of wild type and sex-transformed gonads. A. Heatmap representing significantly differentially expressed genes (p-value<0.05) in XX, XY; *tra*^F^, XY and XX; *tra*^-^ larval gonads by two-way ANOVA analysis of bulk RNA-seq data B. The summary of ANOVA analysis for XX, XY; *tra*^F^, XY and XX; *tra*^-^ gonads representing the number of genes belonging to the Karyotype, TRA only or their Interaction categories. C. Representing encoded Dsx isoforms in different genotypes. XX encodes DSXF and *dsx^D^/dsx^1^* and XX*; tra^-^* encode DSXM. The summary of ANOVA analysis for XX, XX*; dsx^D^/dsx^1^* and XX*; tra^-^* gonads represent the number of genes in *dsx* only, *tra* only or ‘*dsx* by *tra*’ categories. XX; *dsx*^D^ (dsx[1]/dsx(D)) is a progeny from w^1118^; dsx[1] p[p]/TM6 females and w^1118^/Y; dsx(D),Sb1,e1.1(3)e1/TM6 males. D. Splicing analysis comparing XX*; dsx^D^/dsx^1^* and XX*; tra^-^* representing all significantly differential splicing events of TRA (Percent spliced in PSI with log10 Adj.P-value) E. The gene expression (Average Transcript per Million (TPM)) of *nos, nsl-1, wwox Npc1a* in different cell types of larval ovary and testis generated utilizing the scRNASeq.

To better understand the contributions of the two primary sex determination factors (Tra presence and X chromosome number) and the interactions between them, we looked at the sets and intersections of the genes identified from our two-way ANOVA analysis more closely (**Fig. 3B**). As might be expected given the differences in the morphologies of the four gonads types, there were enormous numbers of genes (10,972 genes; ∼80% of the genome) that were differentially expressed (at padj <0.05) in at least one gonad type. Only a few genes were detectably expressed but failed to show differential expression (2,683 genes). We found 1,562 genes whose expression correlated with Tra presence and 777 genes whose expression correlated with X chromosome number. The largest class of 5,174 genes showed expression correlated with Tra presence, X chromosome number, and their interactions. Thus, there were only a few genes that responded cleanly to the primary factors alone. These data suggest a complex set of regulatory pathways, some of which are *tra* regulated, some of which are Tra-independent, but X karyotype-dependent, and a larger set integrating these primary signals.

The gene expression changes in third instar gonads (see **Fig. 1C, E**), due to *tra* could be *dsx* mediated. However, given that *tra* and *dsx* have non-overlapping expression in third instar gonads, it is possible, or perhaps even likely, that there are effects of *tra* that are not mediated by *dsx*. We performed RNA-seq on XX*; dsx^D^/dsx^1^*ovaries transformed into testes in quadruplicates and compared those to the XX*; tra^-^* ovaries transformed into testes. Both encode only Dsx^M^, but through different mechanisms, one via *tra* and another via *dsx* which is downsteam of Tra regulation. On the other hand, XX wild-type ovaries express Dsx^F^. A two-way ANOVA of these XX expression profiles considering the factors *tra* and *dsx* (**Fig. 3C**) revealed 146 genes regulated by *tra*, 5852 by *dsx* and 520 genes by *dsx* by *tra*. First, this suggests that the majority of the effect of *tra* is *dsx* mediated. Second, this suggests considerable *dsx*-independent control of gene expression, but these experiments were done on different genetic backgrounds. This level of differential expression could easily occur due to the heterozygosity for a Deficiency (Df) in the *tra* flies and Dfs are known to result in expression changes (Lee et al. 2018). The presence of two dominant genetic markers [The *Kinked* allele (*Ki*) (Garcia-Bellido and Dapena 1974) is expressed in the somatic cells of the gonads by scRNA-seq, the *Stubble* (*Sb*) (Lindsley and Zimm 2012) allele is unmapped to the genome], and/or other elements of background could also contribute to differential expression in our experiments (**Fig. 3C**).

There is a way to filter for genes that are potentially direct Tra target genes. The above is a gene-level analysis, but genes directly regulated by Tra are expected to differ by splicing pattern, which might also help winnow out effects of background. The gonads show extreme levels of alternative splicing (Chen et al. 2014; Gibilisco et al. 2016), so one cannot simply compare gene expression among wild-type and sex-transformed gonads. Additionally, Dsx binds to many genes implicated in splicing (Clough and Oliver 2012), such that pre-mRNA splicing pattern alone cannot be used to implicate direct action by Tra. To help parse alternatively spliced transcripts regulated by Tra from those regulated by Dsx, we compared RNA-seq profiles of *dsx^D^/dsx^1^*ovaries transformed into testes to those of XX*; tra^-^* ovaries transformed into testes (**Fig. 3D**). We did this using a splicing analysis tool, rMATS (Shen et al. 2014), that detects differential alternative splicing events from RNA-Seq data by comparing exon inclusion levels between two or more sample groups (**Table S6**). We observed only eleven novel significant differential splicing events due to Tra and not Dsx in these experiments. One of these genes producing these Tra-dependent alternative splicing events, *nanos* (*nos*), is highly expressed only in germ cells in the scRNA-seq data (**Fig. 3E**) (Forbes and Lehmann 1998), raising the possibility that Tra functions in the germline, even though it is clearly not required for female fertility. The *non-specific lethal 1* (*nsl-1*) gene encodes histone H4 acetyltransferase complex member (Mendjan et al. 2006; Sheikh, Guhathakurta, and Akhtar 2019) and is also highly expressed in the germline and has Tra-dependent alternative splicing. Several of the Tra-regulated splicing events occurred in transcripts of genes highly expressed in the terminal filaments of the female germ cell niche (Eliazer and Buszczak 2011), where *tra* was highly expressed but *dsx* was not. These include the tumor suppressor homolog *WW domain containing oxidoreductase* (*wwox*), which encodes a putative short-chain dehydrogenases/reductase involved in redox balance and steroid metabolism (Brumby and Richardson 2005; O’Keefe et al. 2011), and *Niemann-Pick type C-1a* (*Npc1a*), which encodes a cholesterol trafficking protein (Huang et al. 2005; Bialistoky et al. 2019). This is exciting given the role of steroid ecdysone in metamorphosis and sex differences in adults (Carney and Bender 2000; Clough et al. 2007; Ables et al. 2016; Grmai et al. 2023). The role of steroids in larval gonad development during the ecdysone pulse is underexplored. Determining if any of these genes are direct Tra targets in the gonads will require much more focused work.

### Germline cell divisions is altered

Sex-transformed gonads exhibit distinct phenotypes due to gene expression changes, which provide insights into biological mechanisms. We observed that Gene Ontology terms for cell division were enriched among the genes expressed in XX ovaries and XY testes transformed into ovaries (**Fig. S2, Table S5**), suggesting that both ovary types have actively dividing cells. For example, the *doubleparked* (*dup*) gene, which is required at origins of replication (Whittaker, Royzman, and Orr-Weaver 2000), was overexpressed in the two types of ovaries (**Fig. 4A**). Single-cell RNA sequencing (scRNA-seq) data on larval gonads show that *dup* is highly expressed in the germ cells and at lower levels in somatic cells (**Fig. 4B, C**). Multiple additional proteins required at the origin of replication, as well as the RNA primase and DNA polymerases, showed a similar pattern (**Fig. S4**). These data suggest increased S-phase activity in XX ovaries and in XY testes transformed into ovaries. Mitotic divisions in these gonads are expected to be of two types: those regenerating the stem cell population and those resulting in the eventual formation of differentiated 16-cell cysts. Differentiation requires the action of *bag-of-marbles* (*bam*) (McKearin and Ohlstein 1995; Gilboa and Lehmann 2004b). We observed much higher levels of germline-limited *bam* in the XY testes transformed into ovaries (**Fig. 4D-F**) than in XX ovaries, suggesting that the greater numbers of germ cells in these sex-transformed gonads could be due to precocious differentiation of germ cell cysts.

**Fig 4.**
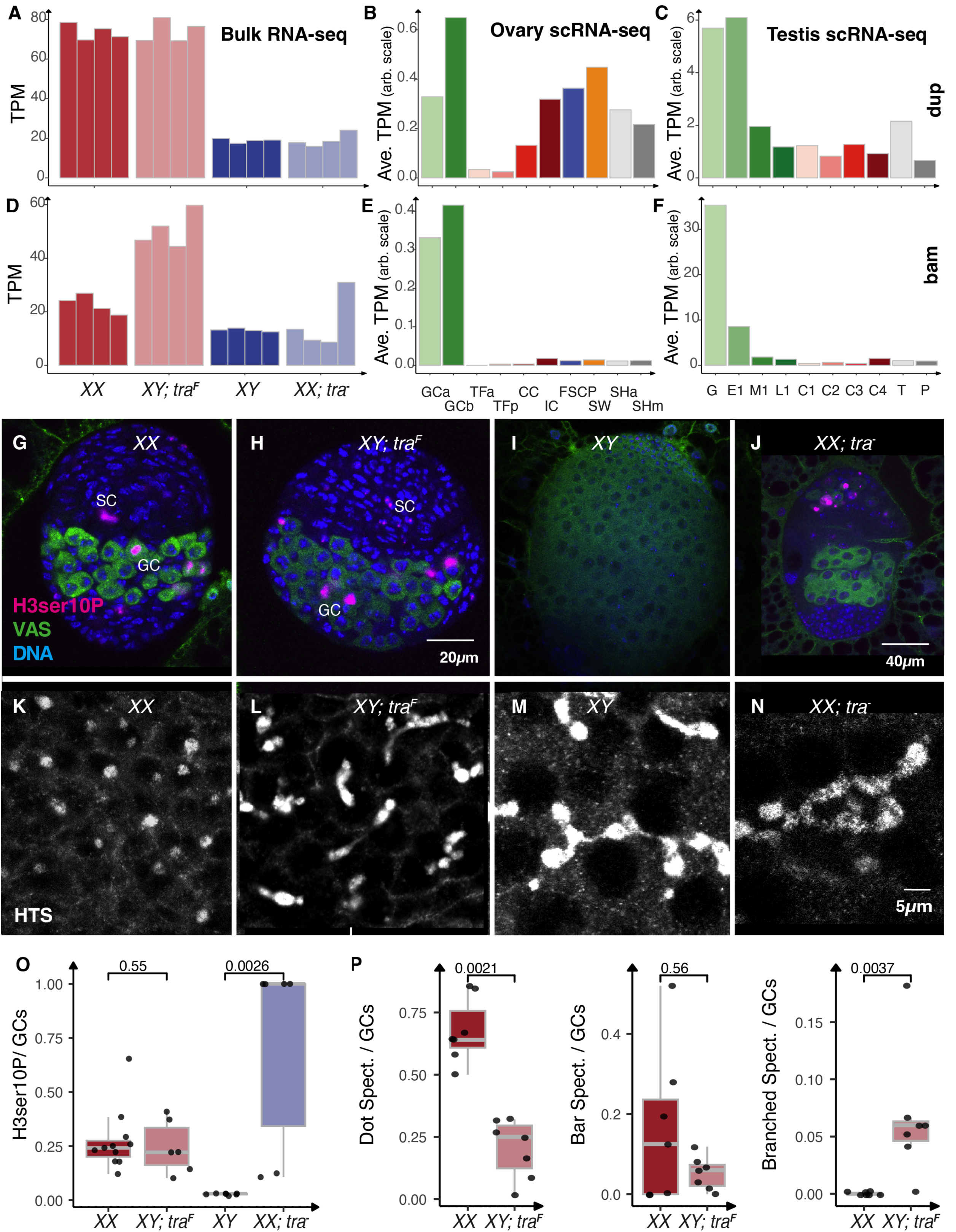
Germ line cell division in the sex-transformed gonads. A - D., The gene expression (Transcript per Million (TPM) or Average TPM) of *dup* and *bam* in bulk RNAseq (from XX, XY; *tra*^F^, XY and XX; *tra*^-^ gonads) and in different cell types of wild type larval ovary and testis by scRNASeq G. - N., Immunostained XX, XY; *tra*^F^, XY and XX; *tra*^-^ gonads to identify dot, bar spectrosomes, and branched fusomes in Germ cells. VAS: green (Germ cells), H3ser10P: Magenta (marks entry into metaphase, mitotically active germ cells,), DAPI: blue (Nuclei), HTS; gray (spectrosome) Scale bar 20 *μ*m in G and H, 40 *μ*m in I and J, 5 *μ*m in K - N. O. The ratio of the number of H3ser10P positive cells to the number of total germ cells in a gonad. P. The ratio of the number of Dot, bar spectrosomes, and branched fusomes to the total number of germ cells in a gonad. Two-sample Wilcoxon Rank Sum (Mann-Whitney) test is utilized to find the statistical significance in O - P

To test the predictions on the cell division and differentiation as the basis of the increased numbers of germ cells in XY testes transformed into ovaries, we stained for Histone H3, Ser10 phosphorylation (H3ser10P), which marks entry into metaphase (**Fig. 4G-J, O**) (Prigent and Dimitrov 2003). We observed mitotically active germ cells in all four genotypes, although the massive numbers of post-mitotic 16-cell cysts in the XY testes resulted in a lower frequency. In the two ovary types, we also observed clear cases of mitotic somatic cells in addition to the germline. Germline stem cell divisions are restricted to the niche, but the divisions we observed were not restricted to those germ cells closest to the niche. Surprisingly, the rates of mitosis did not significantly differ between XX ovaries and XY testes transformed into ovaries (**Fig. 4O**). Given the greater number of germ cells in XY ovaries, these data suggest that some initial increase in germ cell number occurred before the late third instar. We will need to examine earlier developmental stages to explore when this precocious division begins.

Germ cell divisions that regenerate the stem cells undergo complete cytokinesis, whereas divisions that create 16-cell cysts do not. The incomplete cytokinesis in differentiating germline cysts is marked by the presence of cytoplasmic bridges with spectrin-containing bar shaped and later branched fusomes (Lin, Yue, and Spradling 1994; M. de Cuevas, Lee, and Spradling 1996; Petrella, Smith-Leiker, and Cooley 2007). Complete cell division results in dot spectrosomes. We observed many more dot spectrosomes in the XX ovaries and many more bar shaped fusomes in XY testes transformed into ovaries (**Fig. 4K-N, P**). This provides additional evidence of precocious germline development of XY germ cells in the ovary. Both XY testes and XX ovaries transformed into testes were characterized by abundant fusomes (**Fig. 4K-N, P**). XY germ cells in both wild-type testes and in testes transformed into ovaries have begun the amplification divisions characteristic of differentiation.

### Determining the sex of transformed gonads is not straight forward

The XY germ cell mitotic divisions in an ovary could be simply due to precocious or accelerated female germ cell division, or a sex transformation towards male. We know that Sxl protein is required in germ cell cysts of XX ovaries, but not in XY testes. If XY germ cells in an ovary express female *Sxl* mRNAs, then they are likely female. If XY germ cells in an ovary are following a male germ cell differentiation pattern, they might not express female *Sxl* mRNAs. Transcriptional control of Sxl has not been widely studied. We first examined bulk expression of *Sxl* in the sex-transformed gonads, and found the highest levels in XX ovaries, very low levels in XY testes and intermediary levels in the sex-transformed gonads (**Fig. 5A**). In wild-type larval gonads, scRNA-seq data revealed that *Sxl* is most abundant in the germ cells (**Fig. 5A**). These novel data raise the possibility that transcriptional control of Sxl in the germline depends on both the X chromosome karyotype and Tra. It is unclear if this is a specific feedback loop, where Sxl regulates *tra* splicing and Tra regulates *Sxl* transcription, or an indirect effect via Dsx in the soma. In any case, it is clear that *Sxl* transcript levels, not just splicing, are regulated in gonad development.

**Fig 5.**
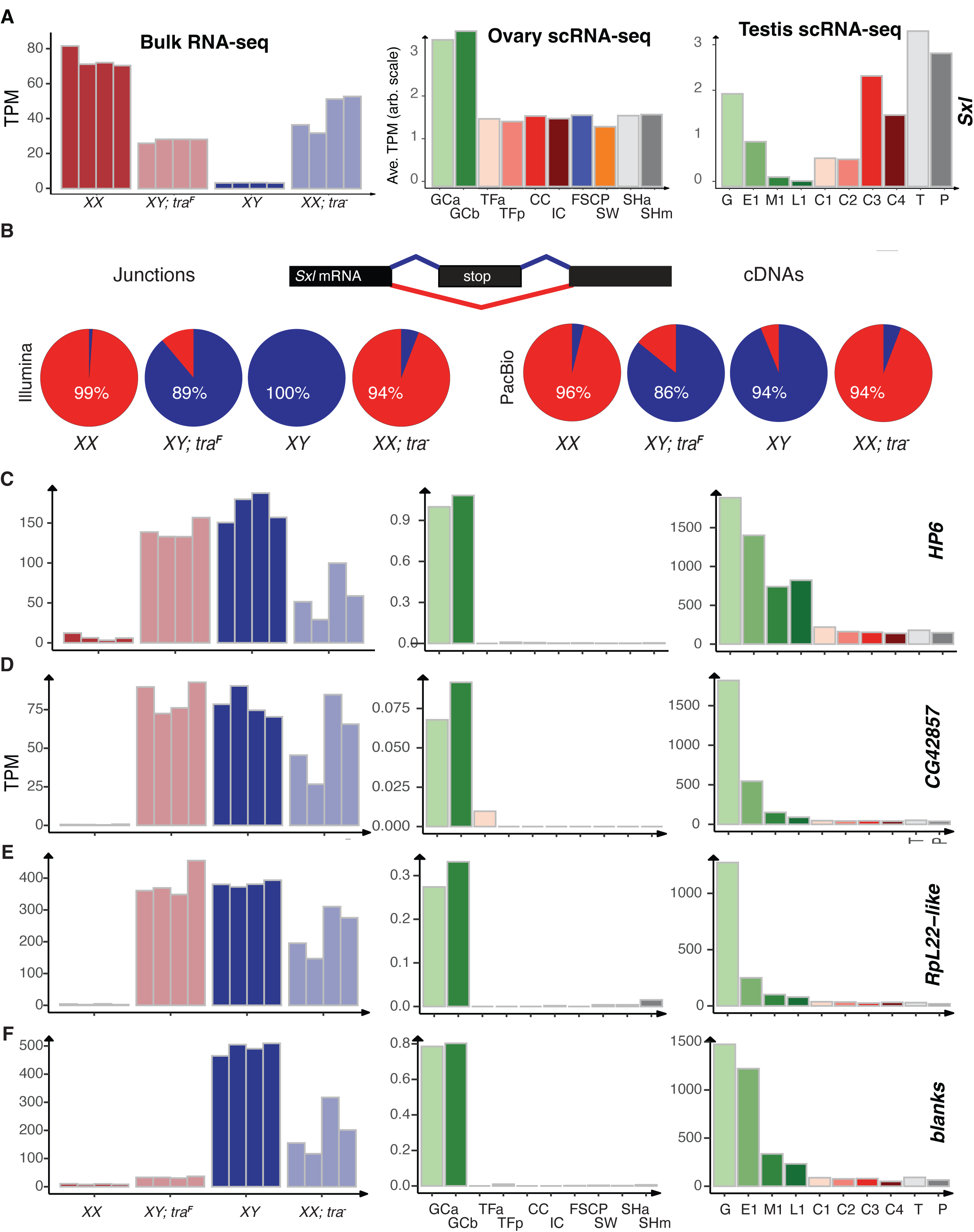
Sexual identity of germ cells in the sex-transformed gonads. A. The *sxl* expression (Transcript per Million (TPM) or Average TPM) in bulk RNAseq and in different cell types of larval ovary and testis by scRNASeq B. Splicing pattern of *sxl* (Percent spliced) in XX, XY; *tra*^F^, XY and XX; *tra*^-^ larval gonads analyzed from Illumina (short reads) and PacBio (long read) RNA-sequencing data C. - F., The gene expression (TPM) of *Hp6, CG42857, RpL22-like* and *blanks* in bulk RNAseq and in different cell types of larval ovary and testis by scRNASeq

Post-transcriptional control of *Sxl* is critical for function. *Sxl* pre-mRNA is spliced into the female isoform that encodes functional Sxl or the male isoform that does not (Cline and Meyer 1996). We examined the splicing patterns of the gonads using both the Illumina junction reads we have already described in this manuscript and a deep PacBio long-read dataset from the same four larval gonad types (**Fig. 5B**). We observed primarily female *Sxl* short read junctions and PacBio cDNAs in RNAs from XX ovaries and XX testes transformed into ovaries. Similarly, we observed more *Sxl* male splice junction reads and cDNAs in the XY gonads. This was consistent with the idea that Sxl pre-mRNA splicing is regulated by germline X chromosome karyotype (Schüpbach 1985; Waterbury et al. 2000; Chau, Kulnane, and Salz 2009; Bhaskar et al. 2022), but it appears to not be a clean on/off switch as the effect of Tra is evident at both the transcriptional and splicing levels. Nevertheless, analysis of *Sxl* splicing, at face value, suggests that XX germ cells are feminized and XY germ cells are masculinized. If the transcript levels are important, then both the XX karyotype and a female *tra* activity are important. These somewhat messy conclusions might help explain why there is a dispute in the literature on what controls Sxl activity in the germline. As we discussed earlier, very few genes respond only to the XX versus XY karyotype. Our data does not support a simple model where Sxl responds to the XX karyotype alone (Grmai, Pozmanter, and Doren 2022).

The Sxl pattern alone does not help us determine if XY germ cells in testes transformed into ovaries and XX germ cells in ovaries transformed into testes are “female” or “male”. If one looks at a larger set of genes, we find examples where the expression of genes in XX germ cells of the testis suggests male identity (**Fig. 5C-F**). For example, the gene encoding Heterochromatin protein 6 (Hp6), which binds telomeres, *CG42857*, *Ribosomal protein L22-like* (*RpL22-like*) (Shigenobu et al. 2006; Kearse, Chen, and Ware 2011), and the siRNA binding protein encoded by *blanks* (Gerbasi et al. 2011) are all highly expressed in XY testes, and scRNA-seq shows that they are restricted to the germline. They are also expressed in XX ovaries transformed into testes. Of these, *Hp6*, *CG42857*, and *RpL22-like* are highly expressed in XY testes transformed into ovaries. At face value, this indicates that XY germ cells remain male in a female ovary. However, *blanks* is very poorly expressed in XY germ cells of testes transformed into ovaries, consistent with the idea that testes development promotes, or ovarian development blocks *blanks* expression. The best resolution to this conundrum is to hypothesize that there are multiple pathways controlling germline sexual identity: some that are based on autonomous germline signals, some rely on sexual signals from the soma, and some that use both.

Examining the expression of genes responding to the presence of *tra* and/or an XX karyotype supports the idea that there is not a clean sexual identity in the germ cells of sex-transformed gonads. The conclusion on sexual identity depends entirely on which genes you explore. We extended this analysis of the RNA-seq data and generated a few new reporter genes to visualize the expression patterns in wild-type gonads. The *defective chorion* (*dec-1*) gene controls the production of the eggshell in adult females (Noguerón, Mauzy-Melitz, and Waring 2000) and we find that it is highly expressed in both XX and XY ovaries, suggesting that it is regulated by Tra only (**Fig. 6A).** In wild-type third instar ovaries, scRNA-Seq datasets showed *dec-1* expression in the intermingled cells surrounding the germline and in the swarm cells, and a *dec-1* reporter we constructed shows a similar pattern (**Fig. 6B, M**). The *dec-1* locus is also expressed in the testes spermatogonia cyst cells (**Fig. 6C, N**), but this expression was in very few cells by microscopy, and was not obvious in the bulk RNA-seq. The *Nascent-associated complex β-subunit-like, testis* 1 (*βNACtes1*) (G. L. Kogan et al. 2017; Galina L. Kogan et al. 2022) gene was expressed in the XY and XX testes, where scRNA-Seq and reporter gene expression showed it was limited to the germline, especially primary spermatocytes (**Fig. 6D-F, N, R**). These data indicate that some aspects of the primary spermatocyte program are executed in XX ovaries transformed into testes. Genes encoding other members of the βNACtes complex (*βNACtes2,3,* and *4*), which associate with the nascent polypeptide in polyribosomes, show the same pattern of expression in scRNA-seq experiments (**Fig. S4**). Interestingly, βNACtes is also germline restricted in embryos, making it an early marker of the germline (Chen et al. 2014). This represents another set of germ line expressed genes that respond to Tra activity, in this case likely by indirect repression. Expression of the histone H4K16 acetyltransferase encoded by *CG1894* was most highly expressed in XY testes and XY testes transformed into ovaries and was expressed specifically in the germline (**Fig. 6G-I, O, S**). This suggests that *CG1894* (S. Zhang et al. 2021) is regulated by X-chromosome number. This is curious, as H4K16 acetylation is associated with X chromosome dosage compensation in the soma (Y. Zhang and Oliver 2007; Kuroda, Hilfiker, and Lucchesi 2016). As a final example, the transcription factor encoded by *ovo* (Pauli, Oliver, and Mahowald 1993; Andrews and Oliver 2002; Bielinska et al. 2005) is highly expressed in XX ovaries, where it is required for germline viability, and weakly expressed in XY testes (B. Oliver et al. 1994; Benner, Muron, and Oliver 2023). The *ovo* locus shows intermediate expression in XY testes transformed into ovaries and XX ovaries transformed into testes (**Fig. 6J**). The *ovo* locus is expressed in XX germ cells in ovaries and in XY early spermatogonia in testes in scRNA-Seq data (**Fig. 6K-L**) and tagged Ovo (Benner, Muron, and Oliver 2023) is found in these same cell types (**Fig. 6P, T**). These data are consistent with reports of *ovo* reporter expression being regulated by both the Tra and X chromosome number (B. Oliver et al. 1994). However, *ovo* is required for germline viability in both XX ovaries and XX ovaries transformed into testes, but not in XY testes nor XY testes transformed into ovaries (Hayashi et al. 2017). The latter indicates that *ovo* is required only in XX germ cells. These are just a few examples showing the diverse responses to mismatched karyotype and sex in the gonads.

**Fig 6.**
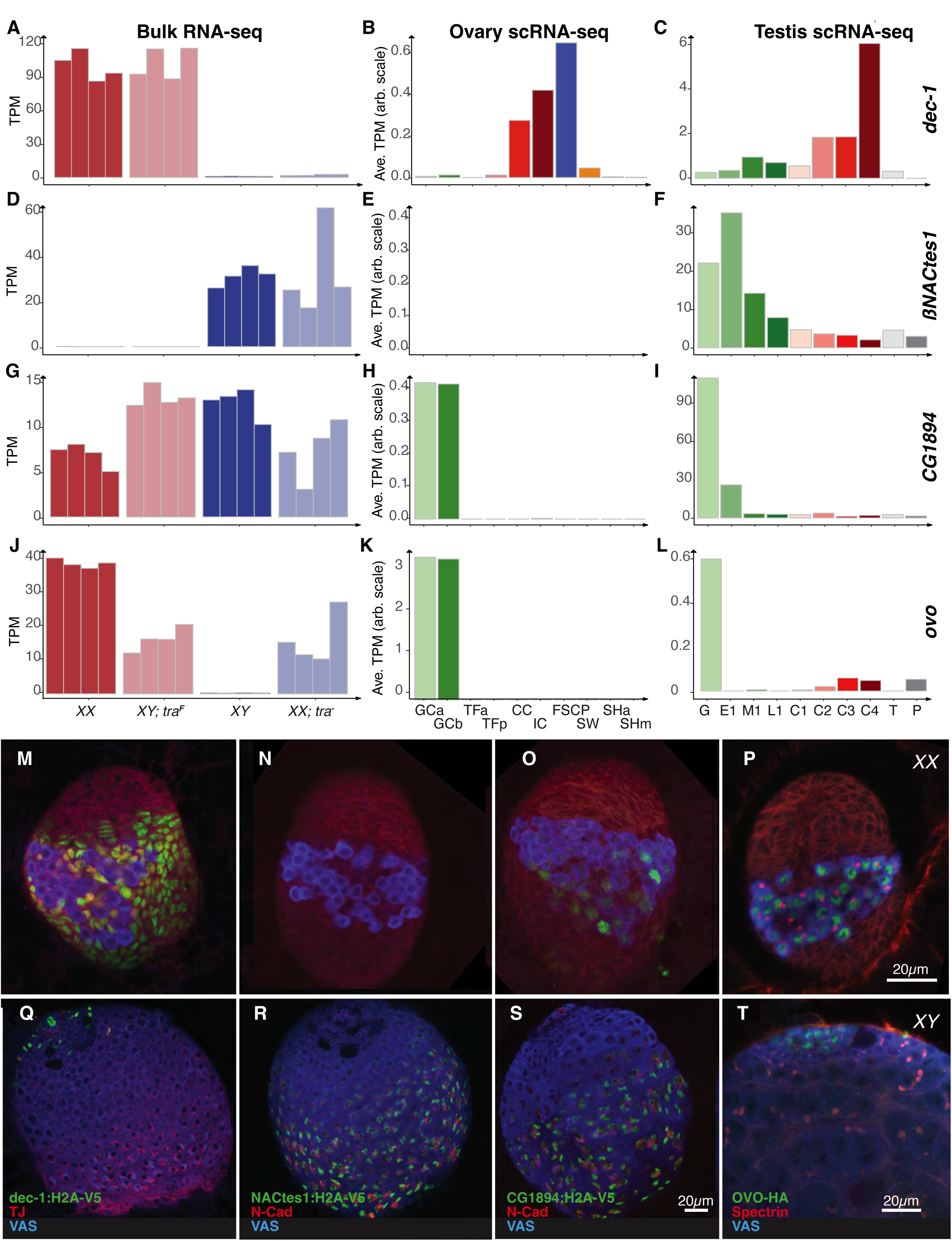
Expression of genes responding to the presence of tra and/or karyotype. A. – L., The gene expression (TPM) of *Ovo, CG1894, bNACtes1* and *dec-1* in bulk RNAseq and in different cell types of larval ovary and testis by scRNASeq M. – T., Immunostained larval ovary and testis to identify different genes. VAS: blue (Germ cells), TJ: Red (cyst cells), N-cad: red (hub), Spectrin; red(fusome), H2A-V5 tagged *Ovo, CG1894, BNACtes1* and *dec-1*: green Scale bar 20 *μ*m

### XX germ cells are not typical Primary Spermatocytes

The primary spermatocyte gene expression profile is remarkably rich. Thousands of genes are highly expressed in these cells and are found nowhere else (White-Cooper 2004). While we did observe expression of a few primary spermatocyte genes (e.g. the *βNACtes* genes discussed earlier) in XX ovaries transformed into testes, the majority of the primary spermatocyte expression profile was highly enriched in XY testes. Indeed, the largest of the clusters in the study are highly enriched for these spermatocyte-specifically expressed genes (**Fig. 3A**, **Fig. 7A**). The reduced expression of these spermatocyte-specific genes in XY testes transformed into ovaries was especially clear, indicating that either XY germ cells cannot enter a normal primary spermatocyte stage in an ovary, or if they do, these cells are rare. However, the analysis of expression in XX ovaries transformed into testes must be tempered due to the highly reduced number of germ cells in these testes. In other words, are these genes reduced in expression because of gene regulation, or the absence of most of the cells of this type? In one of the quadruplicates there is modest expression of some of the early spermatocyte-specifically expressed genes (**Fig. 7A**), suggesting that reduced germ cell number is a factor. This provides more evidence that XX germ cells can at least try to enter spermatocyte stages.

**Fig 7.**
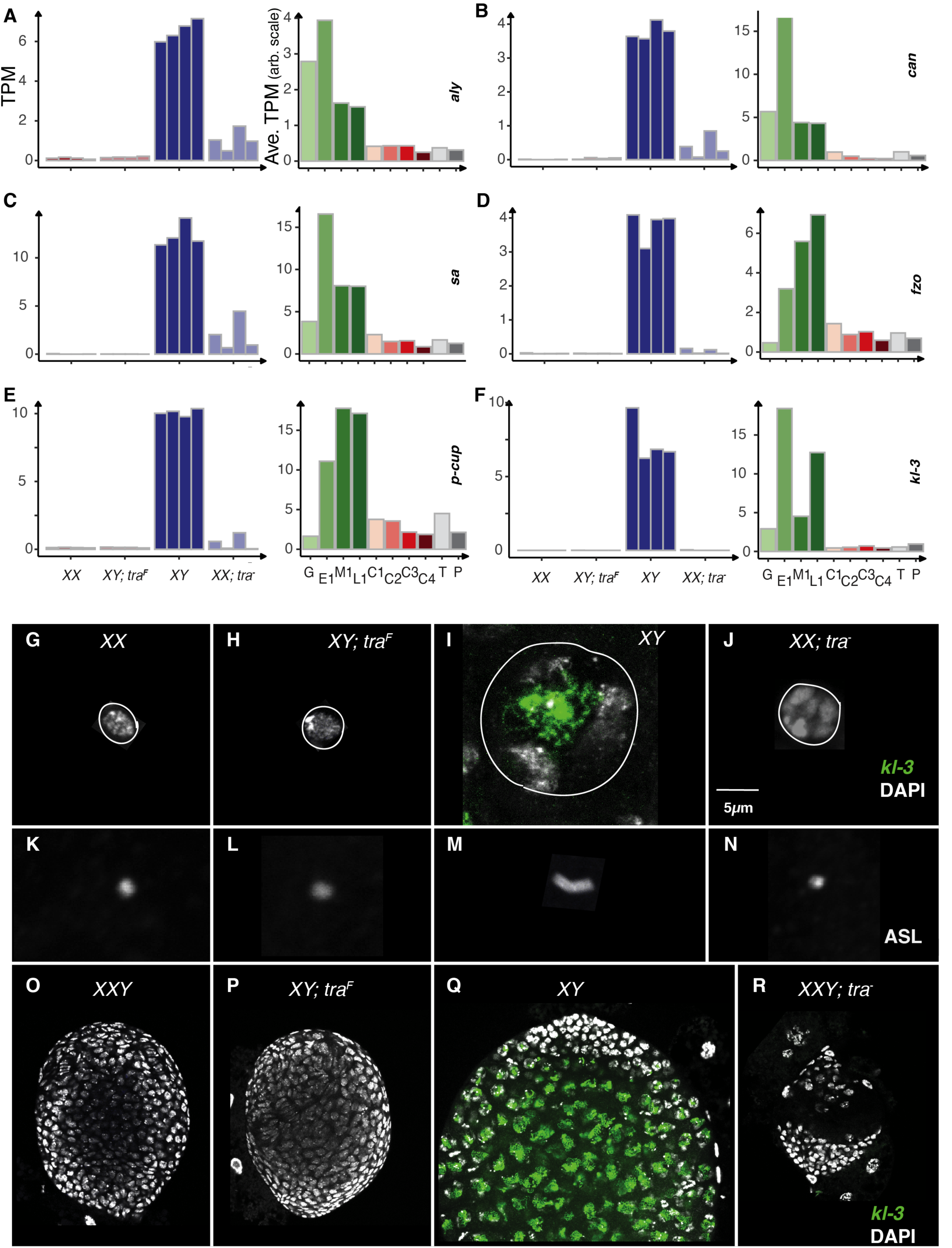
Primary Spermatocytes in sex transformed gonads. A.-F. The gene expression (TPM) of early spermatocyte-specifically expressed genes *can*, *fzo* and *kl-3* in bulk RNAseq and in different cell types of larval ovary and testis by scRNASeq G. - J. Germ line nuclear structure in XX, XY; *tra*^F^, XY and XX; *tra*^-^ larval gonads. DAPI: grey (DNA), *kl-3*: green (*kl-3* transcript in the nucleus). White circle representing the periphery of the nucleus K. - N. Centrioles ASL: grey (Asterless) O. - R. Immuno stained XXY, XY; *tra*^F^, XY and XXY;*tra*^-^ gonads DAPI: grey (DNA), *kl-3*: green (*kl-3* transcript in the nucleus)

There was feeble expression of important initiators of spermatocyte development. For example, primary spermatocytes deploy a specific set of basal RNA polymerase components that are absolutely required for the transition to primary spermatocyte gene expression (Fuller 1998; Lu and Fuller 2015). These include *always early* (*aly*), *cannonball* (*can*), and *spermatocyte arrest* (*sa*) (**Fig. 7A-C)**. These three genes showed highly enriched expression only in XY testes, but the XX ovaries transformed into testes do show weak expression of these genes, while neither XX ovaries nor XY testes transformed into ovaries show evidence of expression (**Fig. 7B**). Genes expressed in primary spermatocytes destined for later spermatogenesis functions are more restricted to XY testes (**Fig. 7D-F**). For example, *fuzzy onions* (*fzo*) encodes an mitofusin utilized in secondary spermatocytes (Hwa et al. 2002), and *presidents-cu*p (*p-cup*) is required for spermatid elongation (Barreau et al. 2008). Weak expression of these genes in XX ovaries transformed into testes also suggests at least an aborted attempt at spermatocyte development. Unsurprisingly, the *male fertility factor kl-3* (*kl-3*) (Pisano, Bonaccorsi, and Gatti 1993; Fingerhut, Moran, and Yamashita 2019a) and other Y-linked genes were not expressed in gonads without a Y (XX ovaries and XX ovaries transformed into testes).

One of the hallmarks of primary spermatocytes is a 20-fold enlargement of these highly transcriptionally active cells (Witt et al. 2019; Raz et al. 2023). The nuclei are also enlarged and show a characteristic chromosome territories phenotype, with the autosomes and X arranged near the periphery and the Y as a diffuse structure in the center (Mahadevaraju et al. 2021). Y-chromosome expression is a quintessential primary spermatocyte activity resulting in the formation of Y-chromosome loops due to the transcription of megabase-long genes (Ball et al. 2023). Additionally, the centrioles duplicate, elongate, and are highly persistent in primary spermatocytes (Riparbelli et al. 2020). This is an excellent marker for spermatocyte maturation.

We observed compact germ cell nuclei in XX ovaries and XY testes transformed into ovaries (**Fig. 7G, H**). The wild-type XY spermatocytes showed the typical chromosome territory nuclear structure and *kl-3* transcripts were detected in the central part of the nucleus (**Fig. 7I**). The XX germ cell nuclei of XX ovaries transformed into testes were partially enlarged but showed an atypical morphology with “lumpy” chromatin (**Fig. 7J**). We detected centrioles with anti-Asterless (ASL). Only the primary spermatocytes of XY testes showed duplicated and elongated centrioles (**Fig. 7K-N**). To explore *kl-3* gene expression in individual germ cells in all wild-type and sex-transformed gonads, we generated gonads where all four of the test genotypes included a Y-chromosome and performed in situ hybridization for *kl-3* expression. We observed kl-3 only in the primary spermatocytes of XY testes (**Fig. 7O-R**). These data suggest that Y-chromosome expression requires a Y-chromosome, a single X chromosome, and a male soma.

## DISCUSSION

We have long known that the sex of the germline and soma must match to ensure gametogenesis in Drosophila (Murray, Yang, and Van Doren 2010). Determining what happens to germ cells placed in a sexually inappropriate environment has been far from clear. In the legacy literature, understanding whether germ cells attempt to follow the developmental program of the rest of the fly or their own internal program has been hindered by the lack of markers. Using the few sex differentiation markers that were available did not result in a convincing determination of the fate of sex-transformed germ cells in adults, and worse, the reports were often contradictory (Brian Oliver 2002). XX germ cells in ovaries transformed into testes were reported to enter either spermatogenesis or oogenesis. Similarly, XY germ cells in testes transformed into ovaries resulted in actively mitotic germ cell tumors that have been interpreted as germline neoplasias or either oogenic or spermatogenic cells that fail to differentiate (Brown and King 1961; Seidel 1963; Nöthiger et al. 1989; Casper and Van Doren 2006).

Examining larval rather than adult gonads has helped us understand more about the nature of the sex-reversed gonad phenotype (**Fig. 8A**). In particular, the adult XX germ cells in ovaries transformed into testes are extremely sparse if present at all (Nöthiger et al. 1989). In larvae, we found that the XX germline stem cells ovaries transformed into testes are often not attached to the hub. Attachment of male germline stem cells to the hub is known to be required for maintaining the stem cell population (Tulina and Matunis 2001), strongly suggesting that a major reason for germ cell absence in these sex-transformed testes is stem cell depletion. Stem cell niche communication involves intimate cell-cell contact using adhesion molecules and ligand/receptor systems (Voog, D’Alterio, and Jones 2008; Marthiens et al. 2010). Sexual mismatches in those systems could easily disrupt contact. Stem cell depletion alone would result in a burst of spermatogenesis in the germ cells detached from the hub (Margaret de Cuevas and Matunis 2011). While we saw limited evidence of some attempted advance to the primary spermatocyte stage, there is clearly more wrong with the germ cells than simple failure of stem cell renewal. The cell morphology lacked the clear subnuclear chromosome territories and did not fully enlarge as occurs in wild-type primary spermatocytes. Thus, in addition to multiple pathways to determine germline sex, there are likely to be different stages of developmental arrest.

**Fig 8.**
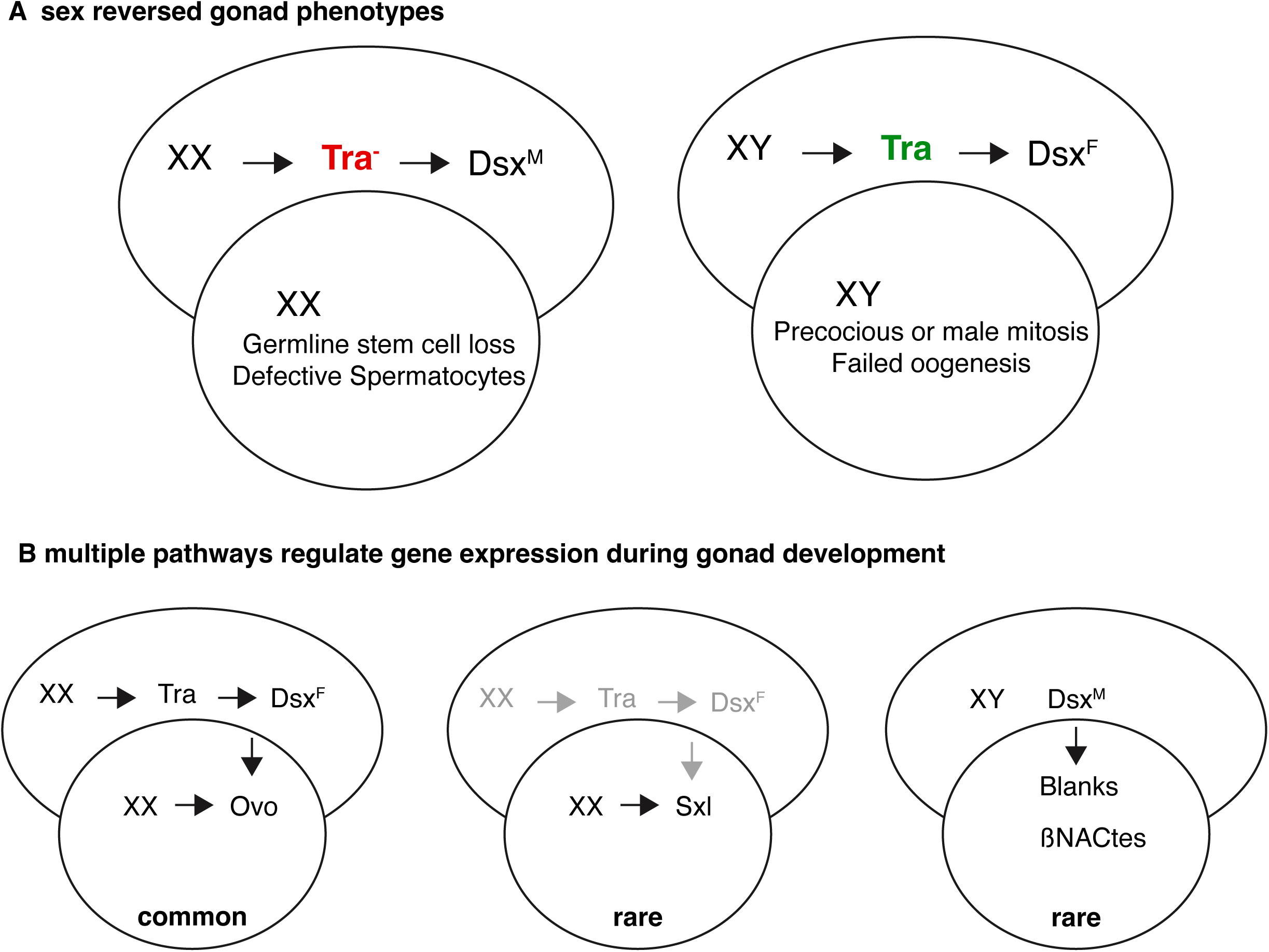
The model explaining the interaction between soma and germ line. A. Nature of the sex-reversed gonad phenotype B. Multiple pathways regulate gene expression during gonad development and controlling sex determination and differentiation of the germline in gonads

The fate of XY germ cells in larval testes transformed into ovaries was less clear (**Fig. 8A**). The major striking morphological differences with wild-type XX ovaries was the presence of extra germ cells. These cells were mitotically active, as were the wild-type controls. Nevertheless, we suggest that the extra numbers of germ cells are due to earlier cell divisions, although we did not determine the timing of this occurrence. In previous work on the male germline niche, activation of XX germ cell divisions can be induced by Stat signaling as occurs in wild-type XY embryos (Tulina and Matunis 2001). The expression of Bam in XY germ cells in testes transformed into ovaries does suggest an attempt at differentiation (Gilboa and Lehmann 2004b).

Because we examined expression in sex-transformed gonads genome-wide, we were able to see multiple patterns suggesting that there is no simple pathway controlling sex determination and differentiation of the germline in gonads **(Fig. 8B)**. In the majority of cases, we found that genes responded both to the XX karyotype and to the Tra status. This includes the *ovo* locus, where previous reporter gene work has shown expression level changes at the germ cell level. We were somewhat surprised to find very few cases where genes with germline restricted patterns were regulated strictly by XX or XY karyotype, and even many of those cases were not clear. Expression of the *Sxl* locus is known to be regulated by X-chromosome counting elements within the germ cells and then autoregulation at the splicing level, suggesting germline autonomous control (Brian Oliver 2002). However, *Sxl* transcript levels clearly responded to Tra status as well in our experiments, and sex-specific splicing was as tightly correlated to X-chromosome number as would be expected in that model. We found a few examples where the sexual differentiation of the testes (in either XY or XX gonads) resulted in clear expression of genes expressed in the male germline. This suggests that there must be a way to bypass the XX karyotype for these genes. In our model we show three examples of pathway configurations, one through the somatic signal (*tra* presence/absence), second through the autonomous karyotype (XX/XY) and another through both (interaction of somatic and germline signals) but there could easily be more. Teasing apart these regulatory connections is needed to understand the regulatory connections between the soma and germline.

During our data analysis, we found that feature plots, which include expression bar plots of a gene in bulk samples, cell type-specific expression from prior single-cell studies, and UMAP/tSNE plots showing gene expression in individual cells, were very useful. As a byproduct of this work, we created a standalone webpage for dynamic viewing of these feature plots. Access details are provided in the Methods section.

## MATERIALS AND METHODS

### Key resources table

All the reagents used in the current study are listed in ART table **(Table S1)** suggested by Flybase.

### Fly stocks

We grew flies on Fly food A in constant light at 25°C and 60% relative humidity. We collected eggs every 8-16 hr from five mated females and allowed larvae to develop until the late third instar larval stage to prepare samples. We have *w*^1118^ females and males as controls. XX; *tra*^-^ (Df(tra)/tra^1^) is a progeny from w^1118^; Df(3L)st-j7, Ki/TM6 females and w^1118^/Y; tra[1]/TM6 males. To identify the *tra* null progeny, first non TM6 larvae are selected they are either X/X; *tra*^-^ or X/Y; *tra*^-^. The X/X; *tra*^-^ have larger gonad than the wild type XX ovary and smaller than XY wild type (which is identical to X/Y; *tra*^-^ (Df(tra)/tra^1^) testis) that can identify under the stereo microscope even before dissecting the larvae. To be certain, these non TM6 gonads are stained together in all the experiments and separated based on their distinct morphology. XY; *tra*^F^ (U2af-*tra*^F^/+; *tra*^5^) is a progeny from X/X; tra[F.U2af50]/+; tra[5] females and FM7, act-gfp/Y; +/+ males. To identify the XY expressing ectopic Tra, first non-gfp expressing larvae are selected they are either XY; *tra*^F^/+ or XY; *+/+*. The XY; *tra*^F^/+ gonads can be clearly distinguished as they are smaller than XY;+/+ testis and almost similar to the size of a wild type ovary. To minimize the genetic background consequences, all the chromosomes in the Df(3L)st-j7 and tra[1] stocks except the chromosome the mutant alleles are residing, are replaced with *w*^1118^ chromosomes. We could not replace the tra[F.U2af50] stock with *w*^1118^ chromosomes because of unknown reasons.

XX; *dsx*^D^/*dsx*^1^ (dsx[1]/dsx(D)) is a progeny from w^1118^; dsx[1] p[p]/TM6 females and w^1118^/Y; dsx(D),Sb1,e1.1(3)e1/TM6 males. To identify the XX; *dsx*^D^/*dsx*^1^ progeny, first non TM6 larvae are selected. They are either XX; *dsx*^D^/*dsx*^1^ or XY; *dsx*^D^/*dsx*^1^. The XX; *dsx*^D^/*dsx*^1^ have larger gonads than the wild type XX ovary and smaller than XY wild type that can identify under the stereomicroscope even before dissecting the larvae. To be certain, these non TM6 gonads are stained together in all the experiments and separated based on their distinct morphology. The *dsx*^D^ allele has the roo element inserted 48bp 3’ of the female specific splice acceptor site leading to the Dsx^M^ isoform production and *dsx*^1^ allele is spliced correctly but is an amorphic allele produces no functional Dsx protein.

To generate expression reporter lines for *dec1*, *βNACtes1*, and *CG1894*, expression constructs were created by placing basal promoters in front of the coding sequence for histone H2A. Three V5 tag sequences were added to the 3’ end of the H2A sequence. The 1 kb region upstream of the transcription start site for the three genes was inserted in front of the basal promoter in each expression construct. This 1 kb sequence was taken to represent the promoter and enhancer regions for each gene, which could then drive cell type specific expression. Injections of expression constructs were performed by BestGene (Chino Hills, CA). The *ovo-HA* (*y^1^ w^1118^ ovo^Cterm-3xFHA^*) line was obtained from Benner et al. (Benner, Muron, and Oliver 2023). These flies were raised at 25 C on standard cornmeal, sucrose, and yeast meal media (Archon Scientific, Millipore Sigma) under constant light and 65% humidity. Adults were allowed to mate and lay eggs for one day before being transferred to a new vial so that their progeny remained uncrowded. For the *dec1:H2A-V5*, *βNACtes1:H2A-V5*, and *CG1894:H2A-V5* reporter lines, larvae that showed dsRed fluorescence under a dissecting scope were selected for dissection to guarantee that they possessed a copy of the reporter construct.

### Sample preparation

We prepared the gonad samples following the GSE125949 extraction protocol. Briefly, we dissected the 3rd instar immobile larvae prior to anterior spiracle extension and prepupal defecation. Staging was aided by staining the gut contents by feeding the larvae with yeast paste mixed with 0.5 mg/ml Sulforhodamine B sodium salt. For RNA-Seq samples we collected larval gonads without fatbody by enzymatically removing fatbody by treating with 0.075 mg/ml Papain and 0.075 mg/ml Collagenase enzymes. Each sample was collected in 350 μl RLT buffer vortexed quickly and frozen in a dry ice ethanol bath and stored at −80°C until RNA extraction.

### Fluorescent *in situ* hybridization

Fluorescent labelled probe against *kl-3* exon1 was generated and *in situ* hybridization was done following the protocol in the article (Fingerhut, Moran, and Yamashita 2019b).

### Immunostaining

After collecting the gonads, we fixed them in 5.14% formaldehyde in PBTX (PBS with 0.1% triton X) for 15 min at room temp (22° C) on a rotator. After removing the fixative, we rinsed the gonads twice and washed twice in PBTX for 10 min later moved to the blocking solution for at least 30 mins in BBTX (PBTX + 0.5% BSA + 2% Normal goat serum), and incubated the gonads in primary antibody for two nights (36-48 hr) at 4° C in BBTX. After removing the primary antibody solution, we rinsed the gonads twice and washed three times for five minutes twice in PBTX and incubated the gonads with secondary antibodies in BBTX for overnight approximately 24 hr) at 4° C. After removing the secondary antibody solution, we rinsed the gonads twice and washed three times for five minutes and incubated the gonads in DAPI solution (1µg/mL in 1XPBS) for a 30 min incubation period. Later DAPI solution was removed, rinsed the gonads twice and washed twice in PBS for five min. We mounted the gonads onto a microscope slide in Vectashield Mounting Medium or Ultramount Aqueous Permanent Mounting Medium. Images were acquired by a Zeiss confocal microscope (Carl Zeiss LSM 780).

### Image Analysis

Images were initially processed with Zen Black software (Carl Zeiss AG) and then moved into FIJI (Schindelin et al. 2012) for further analysis steps. We used length and width to calculate the Volume of each gonad (V=4/3!abc). In order to measure the length of a gonad, first we chose one of the middle Z-stacks where anterior and posterior ends of the gonad are clearly visible. We drew a line from the anterior tip of the gonad to the posterior tip to measure the length in μm. Similarly, we measured the width except drawing a line perpendicular to the length at the widest part of the gonad. We performed Two sample Welch t-test to find significant differences between two sample types. The number of germ cells in a gonad was counted manually from selecting one middle Z-stack of a gonad. To measure the distance between the niche and the nearby germ cells, we selected one Z-stack where the niche is clearly visible. We drew a line starting from the tip of a specific somatic cell surface in the hub or terminal filament to the nearest germ cell surface and recorded the distance in μm. To measure the mitotically active germ cells, where H3ser10P marks the entry into metaphase, we counted the number of H3ser10P/number of germcells in one gonad. As the dot spectrosomes represent either undifferentiated germ cell or complete cell division and the incomplete cytokinesis in cysts represented by bar shaped and later branched fusomes we counted the number of dot spectrosomes in a gonad and calculated the % Dot spectrosomes per total number of germ cells.

### RNA-seq library preparation and sequencing

We published the RNA-seq library preparation and sequencing (accession series GSE205406). Briefly, PolyA+ selection was done using Dynabeads Oligo(dT) 25, mRNA was fragmented, and RT was performed with the second strand synthesis by incorporating dUTPs. We used barcoded adaptors for indexing the libraries (Qiagen, 98 TruSeq v2 kit barcoded adaptors). We sequenced single-ended 50 bp strategy using a HiSeq Illumina 2500.

### PacBio library preparation and sequencing

We published the RNA-seq library preparation and sequencing (PacBio project PRJNA659550). Briefly, we selected polyA+ RNA, made cDNA libraries using the Clontech SMARTer PCR cDNA Synthesis Kit and the Pacific Bioscience SMRTbell Template Kit following the Pacific Biosciences Iso-Seq protocol for Sequel. We used 250-1000 ng total RNA as input to the Clontech SMARTer PCR cDNA Synthesis Kit. Duplicate reactions were performed for each sample. Ten cycles of PCR were used for 1000 ng input and 12 cycles for 250 ng input. The cycling conditions are as follows: Initial denaturation: 98°C for 30s. 10 cycles at the following temperatures and times: 98°C for 10 seconds, 65°C for 15 seconds, 68°C for 10 minutes and final extension: 68°C for 5 minutes. Two size ranges, <4KB and >4 kb, were selected using PB bead size selection. Each range was sequenced separately. We quantified SMRTbell libraries by Qubit Fluorometric Quantitation and qualified by Bioanalyzer before sequencing. Sequencing was performed on the Sequel using Sequel Sequencing Reagent 3.0 with a run time of 20 hours.

### Illumina RNA-seq data processing

An overview of the RNA-seq data analysis workflow is provided in the supplementary (**Figure S1A**) which is a slightly modified version of the RNA-seq workflow developed by the LCDB core (https://github.com/lcdb/lcdb-wf). Briefly, we de-multiplexed and converted Illumina BCL to FASTQ format. FASTQ files were evaluated for quality control using FastQC v0.11.5 (https://www.bioinformatics.babraham.ac.uk/projects/fastqc/), and raw reads were trimmed for Illumina adapters using Cutadapt v4.7 (Martin 2011) with command line parameters -a AGATCGGAAGAGCACACGTCTGAACTCCAGTCA -q 20 --minimum-length 25. Trimmed reads were aligned to the Flybase reference assembly 6.32 for Drosophila melanogaster using STAR v2.7.0 (Dobin et al. 2013) with command line parameters --outFilterType BySJout -- outFilterMultimapNmax 20 --alignSJoverhangMin 8 --alignSJDBoverhangMin 1 -- outFilterMismatchNmax 999 --outFilterMismatchNoverReadLmax 0.04 --alignIntronMin 20 -- alignIntronMax 1000000 --alignMatesGapMax 1000000 --outSAMunmapped None.

### Illumina RNA-seq quantification and normalization

The reads were pseudo-aligned using Salmon v0.13.1 (Patro et al. 2017) with the command line parameters --libType=A --gcBias --seqBias --validateMappings. The transcript-level Salmon abundances were quantified at the gene level using tximport v1.16.1(Soneson, Love, and Robinson 2016) R package with the parameter countsFromAbundance = ‘lengthScaledTPM’. The values in the count slot returned by tximport were further TMM normalized using edgeR v3.32.1 (Robinson, McCarthy, and Smyth 2010). The counts per million were computed on the TMM normalized values by the edgeR function cpm(log = FALSE). In the rest of our analysis, we tried three additional corrections: 1) logarithm (log) transformation of these cpm values, 2) batch-corrected cpm using SVA v3.38.0 (Leek et al. 2012), and 3) batch-corrected log cpm. As we did not observe any substantial batch effects among the replicates, we report the results using normalized cpm values (i.e., without logarithm transformation and batch correction).

### PacBio data analysis

The results on PacBio RNA-seq data were obtained using the IGV genome browser on a BAM file provided to us by the RefSeq and Gene team lead, Terence Murphy. The BAM file was created by mapping PacBio reads to the RefSeq Drosophila genome available at https://ftp.ncbi.nlm.nih.gov/genomes/all/GCF/000/001/215/GCF_000001215.4_Release_6_plus_ ISO1_MT using minimap2 (H. Li 2018).

### ANOVA analysis

We conducted two separate ANOVA analyses on the normalized counts per million (CPM) values. The first analysis involved 16 samples, each with four biological replicates, representing four distinct genotypes characterized by two factors: karyotype (XX or XY) and the presence or absence of the “tra” factor. Specifically, the values of the two factors for the four genotypes are as follows: 1) XX_control (XX, tra present), 2) XY_traF (XY, tra absent), 3) XY_control (XY, absent), and 4) XX_tra (XX, absent). The two-way ANOVA examined the main effects of karyotype and tra on gene expression, as well as any potential interaction between these factors. A significant main effect of karyotype suggests an overall difference in gene expression between XX and XY individuals, regardless of the presence of “tra”. Similarly, a significant main effect of tra indicates a general influence of this factor on gene expression across both karyotypes. The interaction term is crucial in determining whether the effect of “tra” on gene expression depends on the karyotype, revealing if “tra” has a differential impact on XX versus XY individuals. This analysis provides valuable insights into how these factors shape gene expression and can guide further exploration of underlying biological mechanisms. Each gene underwent this two-way ANOVA separately, and the significance of the three tests (individual effects of karyotype and tra, as well as the interaction) was assessed. The P-values for each test underwent multiple-test correction using the Benjamini-Hochberg method, and adjusted P-values were used to determine significance at a false discovery rate (FDR) of 0.05. Genes were categorized based on the significance of these tests into eight categories, ranging from none of the tests being significant to all three tests being significant. Subsequently, genes in each significant category were further classified by visualizing clustered expression heatmaps and manually selecting an appropriate number of clusters. The Supplementary Table S3 provides details on the number of subcategories and the genes within each category. The genes in the rows of the heatmap in Figure 3A were arranged initially based on ANOVA categories. Within each category, they were sorted according to subcategories, and further ordered based on the best adjusted P-values of the three tests.

In our second analysis, we focused on a subset of 12 samples derived from three XX genotypes: XX, XX_tra and XX_dsxD, each comprising 4 replicates. These samples were characterized by two factors: tra (present or absent) and dsx (with male (DSX^M^) or female (dsx^F)^ isoform). Specifically, the values of the two factors for the 3 genotypes are as follows: 1) XX_control (tra present, dsx^F^), 2) XX_tra (absent, DSX^M^), and 3) XX_dsxD (absent, DSX^M^). To unravel the contribution of dsx in this transformation, whether through a direct mechanism or indirectly mediated by tra, we conducted a two-factor ANOVA. The two-way ANOVA examined the main effects of tra and dsx on gene expression, as well as any potential interaction between these factors. Each gene was examined individually, and their respective P-values were adjusted using the Benjamini-Hochberg method for multiple comparisons. Subsequently, Genes were categorized based on the significance of these (FDR cutoff of 0.05 applied to the adjusted P-values) tests into eight categories, ranging from none of the tests being significant to all three tests being significant.

### Pathway analysis

Gene Ontology for each subgroup of ANOVA groups in the heatmap in Fig 3A was separately performed using the Python package GOATOOLS (Klopfenstein et al. 2018) with SlimGO database, which uses Fisher’s exact test, and setting the significance cutoff of the Benjamini-Hochberg corrected Pvalue at 0.05. The set of all genes expressed in at least one of the 20 samples was used as the background list of genes.

### Single cell testis and ovary data

The single-cell visualization (t-SNE) plots in each of the feature plots were created using the Seurat objects obtained from the authors of the respective publications, ovary (Slaidina et al. 2020) and testis (Mahadevaraju et al. 2021). As the Seurat versions used in the publications were significantly older than the current versions (Seurat v5.0.3 and SeuratObject v5.0.1), we needed to convert them using the UpdateSeuratObject() function. The cell-type-specific average expression for each gene was obtained from the supplementary sections of the two publications. To access the values underlying the barplots in the feature plots, we have reproduced the cell-type-specific average expression values of the genes from our bulk data in Supplementary Table S4.

### Visualization

The heatmap of the Z-score of the replicated expression of the four main genotypes used in the two-way ANOVA, where genes in the rows were stratified by ANOVA groups and subgroups, was created using the ComplexHeatmap v2.15.4 (Gu, Eils, and Schlesner 2016) R package.

The upset plot of all genes belonging to the eight ANOVA groups were created using the ComplexUpset v1.3.3 (Lex et al. 2014) R package.

We created a feature plot for each gene expressed in at least one of the 20 Illumina samples. Each feature plot consists of one bar plot for replicate-wise expression in the bulk RNA-seq data generated for the current publication and two bar plots for the cell type-specific average expression in the single-cell RNA-seq data for ovary (Slaidina et al. 2020) and testis (Mahadevaraju et al. 2021), respectively. Each feature plot additionally shows the t-SNE plots of the single-cell expression for ovary and testis, respectively.

To facilitate dynamic viewing of feature plots for all genes genome-wide, we created a standalone webpage. Access it by unzipping feature_plots.zip from https://doi.org/10.5281/zenodo.12752359 and opening FeaturePlots_v1.html in a browser.

### Splicing Analysis

For our differential splicing analysis, we focused on a subset of eight samples derived from two specific genotypes: XX_tra and XX_dsxD, each comprising four replicates. We used rMATS v4.1.2 (Shen et al. 2014) with the parameters --cstat 0.05 and --tstat 6. The genome index required for running rMATS was prepared using STAR v2.7.11b (Dobin et al. 2013) on the FlyBase r6.32 genome and annotation for Drosophila melanogaster. The volcano plot for the differentially spliced events at an FDR cutoff of 0.05 was created using the R package maser v1.13.1 (https://rdrr.io/bioc/maser).

## Data availability

All the raw data are publicly available through NCBI Gene Expression Omnibus (GEO). Illumina RNA-seq data is available at GEO accession number GSE205406 (BioProject PRJNA844981), and PacBio RNA-seq data is available at BioProject accession PRJNA659550. All the processed data is published with this article as Supplementary Tables S1-S6. Additional intermediate data and R objects used in this study are available as Supplementary Dataset (https://doi.org/10.5281/zenodo.12752359).

## Code availability

All codes used in this study are available from the following Zenodo link: https://doi.org/10.5281/zenodo.12752359.

## Author Contribution

Project conceptualizations, design and interpretations: BO, SM; genomic data generation: SM; genomic data analysis: SM, SP; genomic data analysis design & interpretations: BO, TP, SM, SP; splicing analysis SP, LD, DC; fly genetics and data generation: SM, BM, PB, LB; figure generation: BO, SM, SP; original manuscript writing: SM, BO; methods writing: SM, SP, BM; project resources: BO, TP, PB; manuscript reviewing and editing SM, SP, BO, TP. All authors reviewed the final manuscript and accepted.

## Acknowledgements

We thank previous and current members of the Oliver lab, Przytycka lab and LCDB Informatics team for constructive discussions and suggestions. This research was supported in part by the Intramural Research Program of the National Institutes of Health, USA: The National Institute of Diabetes and Digestive and Kidney Diseases (NIDDK) Grant No. ZIADK015600 and National Library of Medicine (NLM) Grant No. LM200887-17. This research work has received funding from the European Union’s Horizon 2020 Research and Innovation Staff Exchange program under the Marie Skłodowska-Curie grant agreement No. 872539 (Pangaia). This research work is also supported by the grant MIUR 2022YRB97K, Pangenome Informatics: from Theory to Applications (PINC). The work utilized resources from the NIH HPC Biowulf cluster (http://hpc.nih.gov), NCBI and FlyBase databases, and the Bloomington *Drosophila* Stock Center.

## List of supplementary tables and figures

**Table S1: ART table**

**Table S2: Quantification**

**Table S3: Gene Level Metadata**

**Table S4: Single Level Cell Metadata (Testis and Ovary in same sheet)**

**Table S5: Gene Level GO Enrichment analysis**

**Table S6: Splicing Metadata**

**Figure S1: Overview of RNA-seq Analysis**

**Figure S2: GO figure**

**Figure S3: betaNCATS2,3,4 feature plots in a single figure**

